# Carbon dioxide sequestration into biomineral armor by ants

**DOI:** 10.64898/2026.01.21.700952

**Authors:** Hongjie Li, Yihang Fang, Weiqiang Li, Jing Liu, Chang-Yu Sun, Xue Kang, Gaspar Bruner-Montero, Joseph Sardina, Xiaochang Mo, Jia-Long Hao, Jianchu Mo, Lei Cheng, Zhaoming Liu, Ted R. Schultz, Richard E. Johnston, Cameron R. Currie

## Abstract

Over geologic time, Earth’s climate has been shaped by the capture and conversion of atmospheric carbon dioxide (CO_2_) into stable carbonate minerals, including dolomite [CaMg(CO_3_)_2_]. Accelerating natural carbon mineralization offers significant potential for mitigating anthropogenic climate change. Using stable carbon isotope tracking, nano-scale secondary ion mass spectroscopy, and ^13^C SSNMR, we show that, paralleling global biosphere-level processes, *Sericomyrmex amabilis* fungus-farming ants rapidly convert CO_2_ in their nest chambers into a biomineral layer covering their exoskeletons. We further reveal that biogenic carbon mineralization by these ants produces partially ordered dolomite. This rapid sequestration of CO_2_ into defensive armor in ants provides a fascinating natural example of mediation of potentially toxic accumulation of atmospheric CO_2_ that could inform human efforts to mitigate climate change.

---

Atmospheric carbon dioxide (CO_2_) serves as a major reservoir of the global carbon cycle, mediating the exchange of carbon between Earth’s atmosphere and biosphere (*1*). Carbon enters biological systems primarily through the fixation of CO_2_ into organic matter, where photosynthetic organisms, including plants, algae, and cyanobacteria, convert atmospheric CO_2_ into the carbohydrates, lipids, and proteins that fuel terrestrial and aquatic food webs and ultimately support nearly all life on Earth (*2*). In extreme environments, certain chemolithoautotrophic microorganisms fix CO_2_ through non-photosynthetic pathways (*3*). While these organic carbon pathways are essential for sustaining ecosystems, the carbon they capture cycles relatively rapidly back to the atmosphere through respiration, decomposition, and consumption, typically residing in the biosphere for timescales ranging from days to centuries (*4*). The metabolic capacity to fix atmospheric carbon dioxide is unknown in fungi, animals, and most bacteria, underscoring the biochemical challenges of capturing and reducing atmospheric carbon (*5*).

In contrast to the dynamic cycling of organic carbon, the conversion of atmospheric CO_2_ into inorganic carbonate minerals provides the most stable and quantitatively significant long-term carbon storage on Earth, with the vast majority of the planet’s carbon sequestered in carbonate rocks formed over geological timescales (*6*). Carbonate formation occurs through both abiotic and biotic processes that span multiple spatial and temporal scales. Abiotic carbonate precipitation occurs through weathering of silicate rocks, in which atmospheric CO_2_ dissolved in water reacts with calcium and magnesium ions to form the carbonate minerals such as calcite and dolomite. This is a process that operates over millions of years and, by increasing in rate with increasing atmospheric CO_2_ levels, represents a critical negative feedback in Earth’s climate system. Biologically-mediated carbonate mineralization occurs through diverse mechanisms, including the controlled precipitation of calcium carbonate shells and skeletons by marine organisms such as foraminifera, coccolithophores, mollusks, and corals (*7*–*9*), as well as microbially-induced carbonate precipitation in which bacterial metabolic activities alter local chemical conditions to promote mineral formation (*10*). Given the exceptional stability and scale of carbons sequestered as carbonates, accelerating both abiotic and biotic carbonate mineralization for enhanced natural carbon sequestration has emerged as one of the most promising climate mitigation strategies being explored to offset rising anthropogenic CO_2_ emissions (*11, 12*).

One of the most remarkable examples of carbonate biomineralization is the epicuticular high-magnesium calcite armor recently discovered in the leaf-cutter ant *Acromyrmex echinatior* (*13*). *Acromyrmex echinatior* belongs to the ant subtribe Attina, collectively referred to as fungus-farming ants based on their ancient obligate mutualism with specialized fungi that they cultivate for food. Fungus-farming ants forage for vegetative substrates to nourish their fungal cultivars, and in exchange the fungi serve as the primary food source for the ants (*14*). This intensive agricultural lifestyle creates densely populated colonies that represent attractive targets for predators and pathogens. In *Ac. echinatior*, worker ants precipitate a continuous layer of high-magnesium calcite crystals Ca_1.3_Mg_0.7_(CO_3_)_2_ directly onto their exoskeletons through what appears to be a protein-mediated nucleation process (*13*). The biomineral layer, composed of randomly oriented rhombohedral crystals 3-5 μm in size, forms rapidly during worker maturation and increases exoskeletal hardness by more than two-fold despite representing only 7% of total cuticle thickness. This biomineralized armor provides critical protection against aggressive encounters with competing ant species and reduces susceptibility to entomopathogenic fungal infections, indicating that controlled carbonate biomineralization evolved as an adaptive response to ecological pressures faced by agricultural insect societies.

## Spatial and temporal distribution of the ant carbonate minerals

To further explore mineral armor in ants, we investigated the possible presence of biominerals in several other genera of Attina, including *Sericomyrmex. Sericomyrmex* is a genus comprising 11 described species with a distribution from Mexico to southern Brazil, often occurring in clay soils within which they cultivate their fungal mutualist, the primary food source for the colony (Fig. 1A). The genus is distinguished morphologically by its densely pubescent integument covered in short, appressed hairs that give workers a silky or velvety appearance (Fig. 1A, Fig. 2A), from which the generic name derives. 3D reconstruction of *S. amabilis* using X-ray microscopy data reveals that the exoskeletons of workers are variably covered with a granular coating (Fig. 1B), which we report here is an outer layer of crystalline mineral (Fig. 1C). SEM and cryo-SEM images of fractured cuticular cross-sections of *S. amabilis* ants show a distinct interface between the mineral layer and the cuticle (Fig. 1D, fig. S1). Intriguingly, crystals even form in the spaces between the facets (ommatidia) of the compound eyes of mature workers, leaving the centers of the ommatidia free of obstruction, thus apparently conferring increased protection while avoiding loss of vision (Fig. 1E). In young *S. amabilis* workers the biomineral is absent, including around the compound eye (Fig. 1F), but in mature workers the biomineral is distributed across the entire body, covering almost all the cuticle (Fig. 2). Using backscattered electron (BSE) imaging, we reveal that the outer layer is brighter than the cuticle, indicating that it has a higher average atomic number and thus contains heavier elements, and that it covers the entire body, except for the antennae and mandibles (Fig. 2B). In addition, X-ray microscopy analyses of *S. amabilis* ants reveal that the thickness of the biomineral crystals ranges from 7 to 20 μm (Fig. 2C, and movie S1). In general, the dorsal body parts have thicker mineral layers than the ventral parts (figs. S2 and S3, table S1, and movie S2). Notably, several body parts lack mineralization, such as the funiculus (ends of the antenna) and the tarsi (ends of the legs), likely due to their roles in chemoreception and flexible movement (fig. S4).

**Fig. 1.**
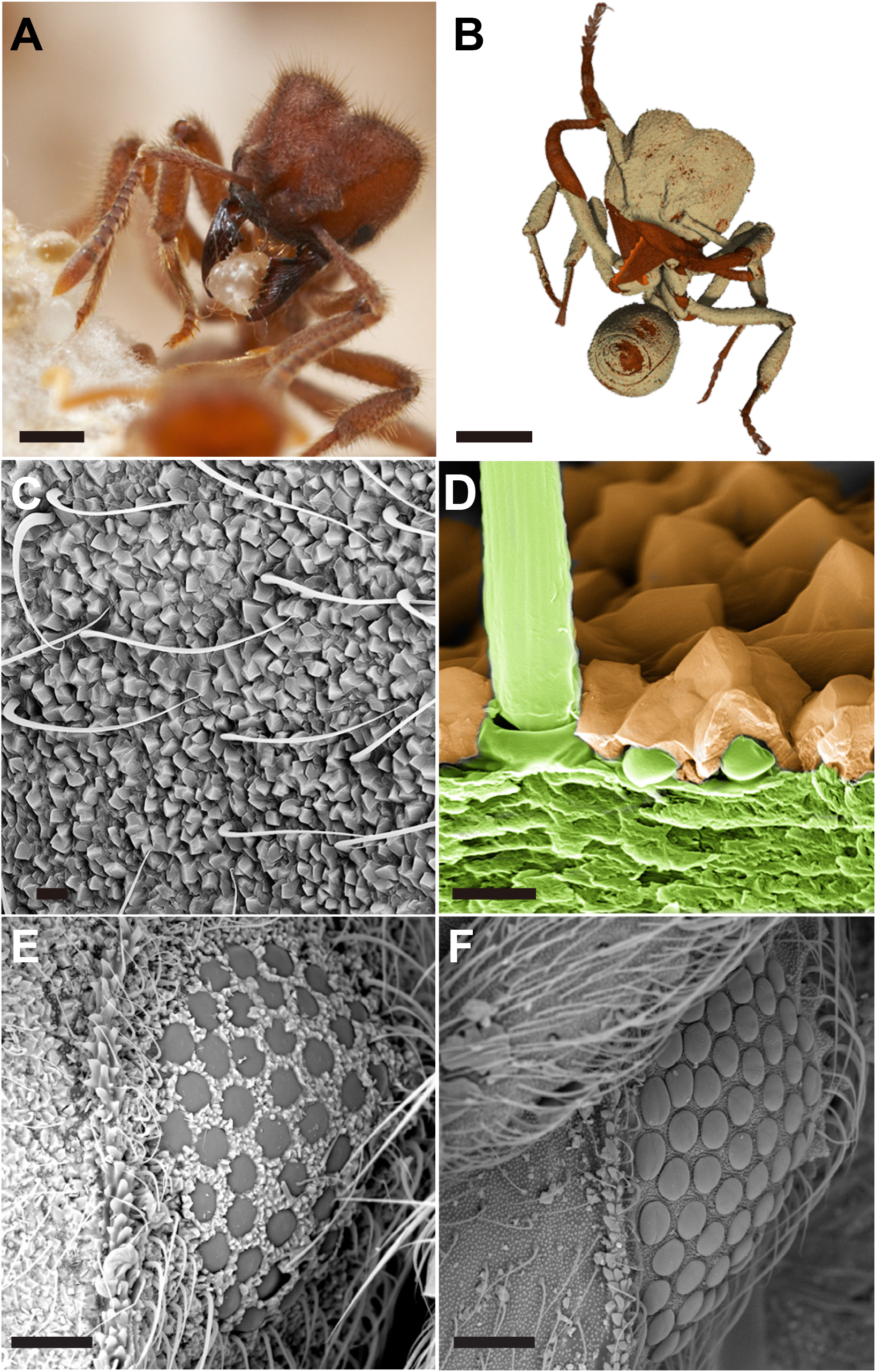
Morphological and structural features of the crystalline mineral layer in the fungus-growing ant *S. amabilis*. (**A**) A worker ant within the colony fungus garden. (**B**) 3D reconstruction of *S. amabilis* using X-ray microscopy showing that a granular coating (falsely colored white) is distributed over the cuticle. (**C**) SEM imaging of dorsal surface of worker ant showing that this granular material is a crystalline layer covering the exoskeleton. (**D**) False-colored SEM image of a fractured cuticular cross-section displaying the distinct organic-mineral interface between the ant cuticle (green) and mineral layer (light brown). (**E** and **F**) SEM images of the ant compound eye, which is nearly covered with crystals in mature (E), but not in young worker ants (F). Scale bars: 1 mm (A and B), 1 μm (C and D), and 10 μm (E and F). [Photo credits: (A) Alex Wild]

**Fig. 2.**
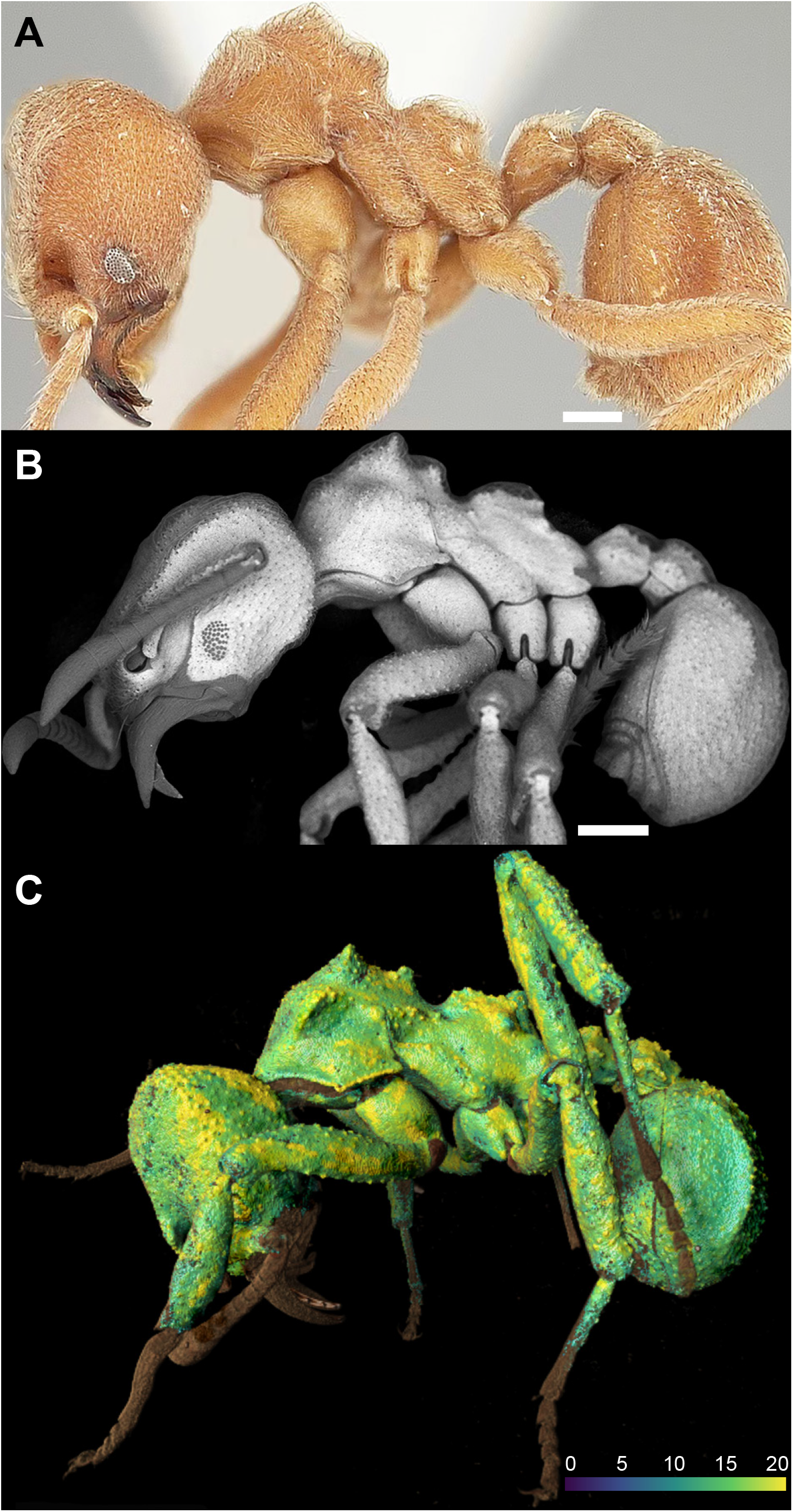
Distribution and thickness of the mineral layer in *Sericomyrmex amabilis*. (**A**) Optical micrograph of a *S. amabilis* worker ant, which have ferruginous brown cuticles that are densely covered by long and flexuous hairs. (**B**) SEM-BSE indicates that the crystalline material is composed of heavier elements (white) relative to the ant cuticle (gray), with the mineral layer covering the entire body of the ant except for the antennae and mandibles. Scale bars: 1 mm (A and B). (**C**) The thickness of the mineral layer ranges from 7 to 20 μm, as measured by X-ray microscopy, with the dorsal regions generally thicker than the ventral regions. Brown color indicates the absence of minerals and the green-to-yellow color scale indicates the thickness of the mineral layer (μm).

## Detection of atmospheric CO_2_ incorporation into the ant carbonate minerals

Discovery of a carbonate biomineral in *Sericomyrmex* is unexpected. Unlike in *A. echinatior*, which until now was the only ant known to form carbonate biominerals, *Sericomyrmex* spp. do not house ectosymbiotic *Pseudonocardia* bacteria on their exoskeletons (*15*). Although the full role of *Pseudonocardia* in the formation of the biomineral in *A. echinatior* remains unclear, workers without *Pseudonocardia* do not exhibit normal biomineral formation, suggesting an essential role for the bacterium, perhaps including carbon fixation. Thus, our finding of calcareous mineral layers in *Sericomyrmex* suggested that these ants may be able to fix carbon without the aid of bacteria, and that the mineral layer could represent a case of a terrestrial metazoan engaging in a close biogenic parallel to geologic carbon mineralization, directly converting atmospheric CO_2_ into a stable carbonate mineral. To explore this possibility, we employed stable isotope tracer experiments in combination with nanoscale secondary ion mass spectrometry (NanoSIMS) and nuclear magnetic resonance (NMR) spectroscopy. First, we conducted *in situ* measurements of carbonate precipitation among different age castes of *Sericomyrmex* worker ants and their fungus gardens. We found that carbonates are present on all mature ants, including on dead workers deposited in the colony refuse mounds where live ants discard dead nest mates, but that carbonates are absent in callow workers and males, and less abundant on gynes (reproductive female alate individuals) and young workers (Fig. 3A, fig. S5). Importantly, they are completely absent from the fungus garden, eliminating the possibility that microbes from within the garden microbiome contribute to the biomineralization. To determine whether atmospheric CO_2_ is at least in part the source of the carbon in the carbonate, we conducted stable isotope ^13^CO_2_-labelling experiments using sub-colonies of *S. amabilis* reared on aliquots of fungus garden (*16*). Sub-colonies of *S. amabilis* were placed in airtight containers in which the internal atmosphere consisted of a mixture of 8% ^13^CO_2_ (purity>99.9%, ^13^C, 99 atom%) and 92% atmospheric air (Fig. 3B). After just 0.5 hours, we observed elevated δ^13^C values [expressed in delta notation: δ^13^C = (R_sample_/R_standard_ − 1) × 1000, R = ^13^C/^12^C, the standard is VPDB] in the ant biomineral, indicating rapid formation of new layers composed of enhanced ^13^C. Across all time points, we observed a consistent trend of increasing CO_2_ fixation with increasing inoculation time, with major ^13^CO_2_ enrichments detected at 10 days (Fig. 3C). Notably, significant enhancements of δ^13^C values were found in the biomineral samples of worker ants, but ^13^C was absent in callow worker extracts (fig. S6), further confirming that the source of the carbon within carbonates is atmospheric CO_2_.

**Fig. 3.**
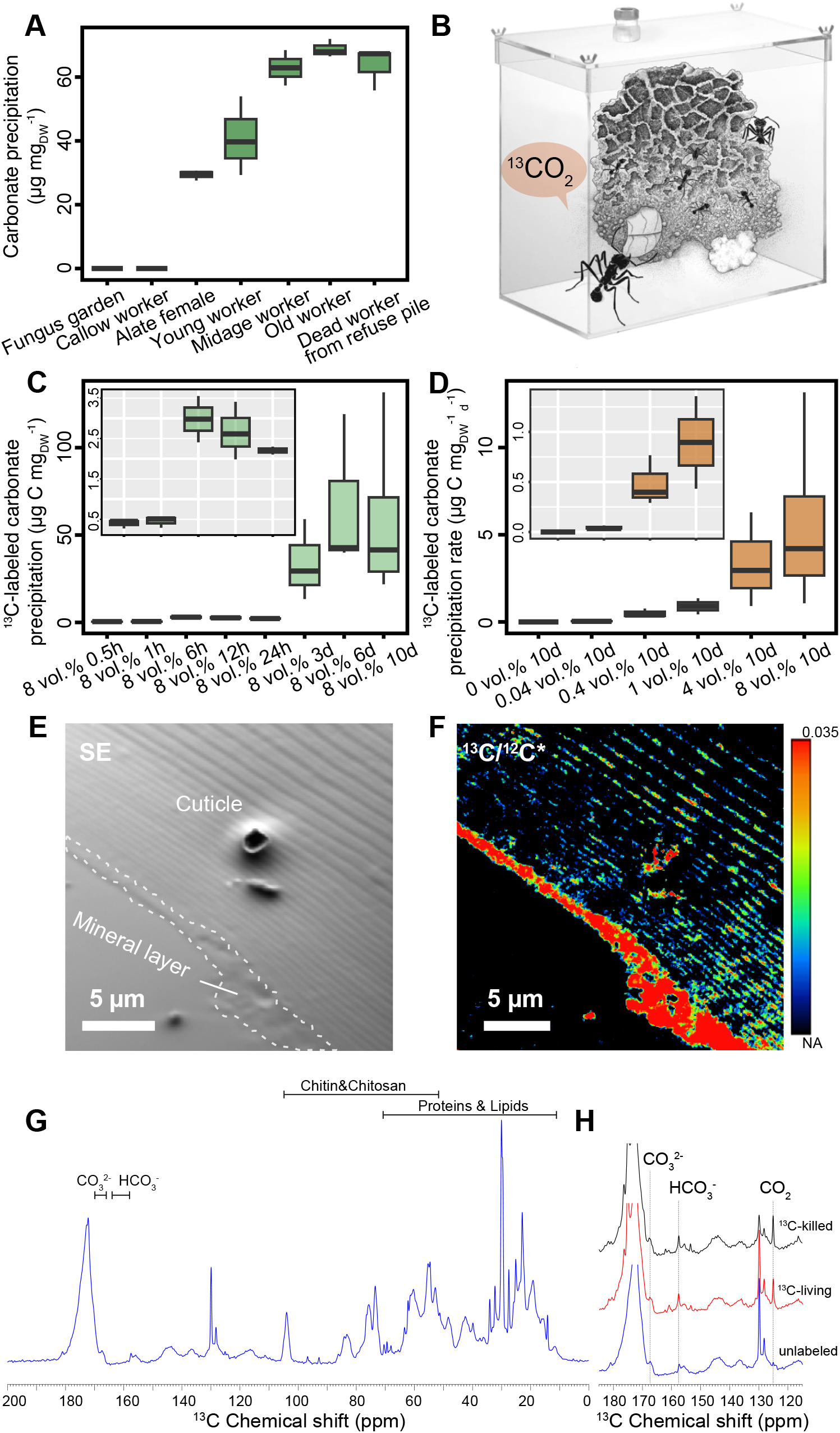
Ant-associated CO_2_ sequestration and distribution of CO_2_ fixation activity. (**A**) *In situ* measurements of carbonate precipitation on different *Sericomyrmex* ant workers of different ages and in their fungus garden. DW, dry weight of ant. **(B)** Depiction of the incubation set-up, enabling the detection of worker ants integrating the ^13^C-enriched CO_2_. **(C)** Measurements of ^13^C in worker ants raised in sub-colonies with an internal atmosphere composed of 8% ^13^CO_2_ for up to 10 days. (inset: magnified plot for the 0.5, 1, 6, 12, and 24 hour-treated samples) **(D)** Quantification of ^13^C in worker ants raised in sub-colonies with varying concentrations of ^13^CO_2_ for ten days. (inset: magnified plot for the 0, 0.04, 0.4, and 1 vol. % samples). Note the precipitation rate is presented as μg ^13^C sequestered per mg dry weight (DW) of ant per day. **(E)** Correlative imaging of the cross-section of ant cuticles and **(F)** the corresponding NanoSIMS image showing ^13^C enrichment. ‘SE’ denotes secondary electron image and * denotes that the ^13^C/^12^C image has been background-subtracted and denoised to enable clearer visualization. NA, natural abundance. **(G)** ^13^C quantitative NMR whole spectrum of unlabeled ant cuticle. **(H)** ^13^C quantitative NMR spectra of ant cuticle before and after ^13^CO_2_ incubation, with 180 to 120 ppm spectra for unlabeled (blue line), living (red line), and heat-killed ants (gray line).

To further quantify CO_2_ conversion into ant carbonate, we analyzed ^13^C incorporation at varying ^13^CO_2_ concentration levels for 10-day periods (Fig. 3D). Elevated ^13^C incorporation was detected in 0.04 vol.% ^13^CO_2_ conditions, indicating that ant carbonate precipitation occurs at normal atmospheric CO_2_ concentrations. Ant-associated ^13^CO_2_ fixation was the highest at 8 vol.% ^13^CO_2_, which mimics CO_2_ levels in the ant nest (*17, 18*). Overall, the ant carbonate ^13^C enrichment was directly correlated with atmospheric ^13^CO_2_ availability, suggesting direct incorporation of atmospheric carbon into the ant carbonate. This was further corroborated by NanoSIMS of cross-sectioned ant cuticle, confirming localized ^13^C incorporation into the ant carbonate mineral layer, with highest ^13^C/^12^C value at 0.035 (Fig. 3, E and F). Next, we performed quantitative analysis of chemical shift peak intensity using one-dimensional (1D) multiple cross-polarization (MultiCP) solid-state nuclear magnetic resonance (NMR) spectroscopy. In ^13^CO_2_-incubated samples from both living ants and dead ant cuticles, we observed enhanced ^13^C resonances, with chemical shift peaks at 125.5 ppm, 157.8 ppm, and 168.7 ppm, corresponding to dissolved carbon dioxide, bicarbonate, and carbonate, respectively (Fig. 3G) (*19, 20*). Our NMR results provide direct evidence that the formation of bicarbonate on the cuticle of *S. amabilis* uses atmospheric CO_2_.

## Detection of near-stoichiometry and partially ordered dolomite in the ant carbonate minerals

Next, we characterized the carbonate mineral layer in *S. amabilis*. High magnification SEM imaging of the mature *S. amabilis* epicuticle revealed clear rhombohedron carbonate crystals (Fig. 4A). Based on bright-field transmission electron microscopy (TEM), selected area electron diffraction (SAED), and TEM-based energy dispersive X-ray spectroscopy (TEM-EDS), the biomineral is carbonate with chemically heterogeneous crystals and with clearly observable Ca-Mg ordering crystal structure that as a result of an ordered alternation of layers of Ca^2+^ and Mg^2+^ ions interspersed with CO_3_^2-^ anion layers (Fig. 4B, figs. S7 and S8). SAED analyses of individual *S. amabilis* epicuticular crystals confirm the dolomite structure, with prominent “a”-reflection (012) and 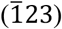, as found in *Ac. echinatior* (*13*), as well as an extra set of super-lattice reflections, such as (101) and 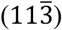 that are absent in calcite structure, which is indicative of Ca-Mg ordering (Fig. 4C, figs. S9, S10, and S11). The observed Ca-Mg ordering indicates that the biomineral in *Sericomyrmex* contains some partially ordered dolomite, a form of biomineral currently unknown in any metazoan. To further corroborate the occurrence of partially ordered dolomite in the biomineral layer, we conducted TEM-EDS, Raman spectroscopy analyses and Attenuated Total Reflectance Fourier Transform Infrared (ATR-FTIR), including samples of geological calcite, geologic dolomite, the spine of the sea urchin *Strongylocentrotus purpuratus*, as well as several other fungus-farming ants. The Raman spectra of *S. amabilis* crystals are consistent with those of geologic dolomite (dolostone), with peaks of lattice mode (L), in-plane bending mode (*v*_4_), and symmetric stretching mode (*v*_1_) shifting toward higher wavenumbers, as compared to both calcite and *Ac. echinatior* and *Apterostigma dentigerum* crystals, the fungus-farming ants with high-magnesium calcite (Fig. 4D, fig. S12). Similar patterns were also observed in the ATR-FTIR spectra (fig. S13, table S2). Powder XRD of the cuticular coating for *S. amabilis* revealed a clear smaller (104) *d*-spacing value of 2.923 Å and detected a super-lattice ordering peak (015) (Fig. 4E, fig. S14, and detailed unit-cell parameters in table S3), both indicative of the presence of partially ordered dolomite (*21*–*23*). Direct measurement of the biomineral layer of *S. amabilis* with quantitative electron probe micro-analysis (EPMA) revealed a magnesium concentration of 42-45 mol.%, which is compositionally similar [Ca_1.1_Mg_0.9_(CO_3_)_2_] to modern geologic dolomite stoichiometry (Fig. 4F, fig. S15). In contrast, the biomineralized armors of the fungus-farming ants *Ac. echinatior* and *Ap. dentigerum* contain high-magnesium calcite [*d*_(104)_ of 2.934 Å, 34.1 ± 1.8 mol.% MgCO_3_; *d*_(104)_ of 2.946 Å, 38.9 ± 0.4 mol.% MgCO_3_, respectively] (table S4, supplementary text for the MgCO_3_ mol.% cut-off of Ca-Mg carbonate terminology). There is a clear trend of increasing hardness with increasing magnesium content across all samples examined (Fig. 4G, fig. S16), with the biomineral layer of *S. amabilis* most closely approaching the hardness levels of geologic dolomite. Notably, enriched Ca and Mg isotope tracer experiments indicated that metal ions within ant biomineral layers derived from their fungal diets (table S5).

**Fig. 4.**
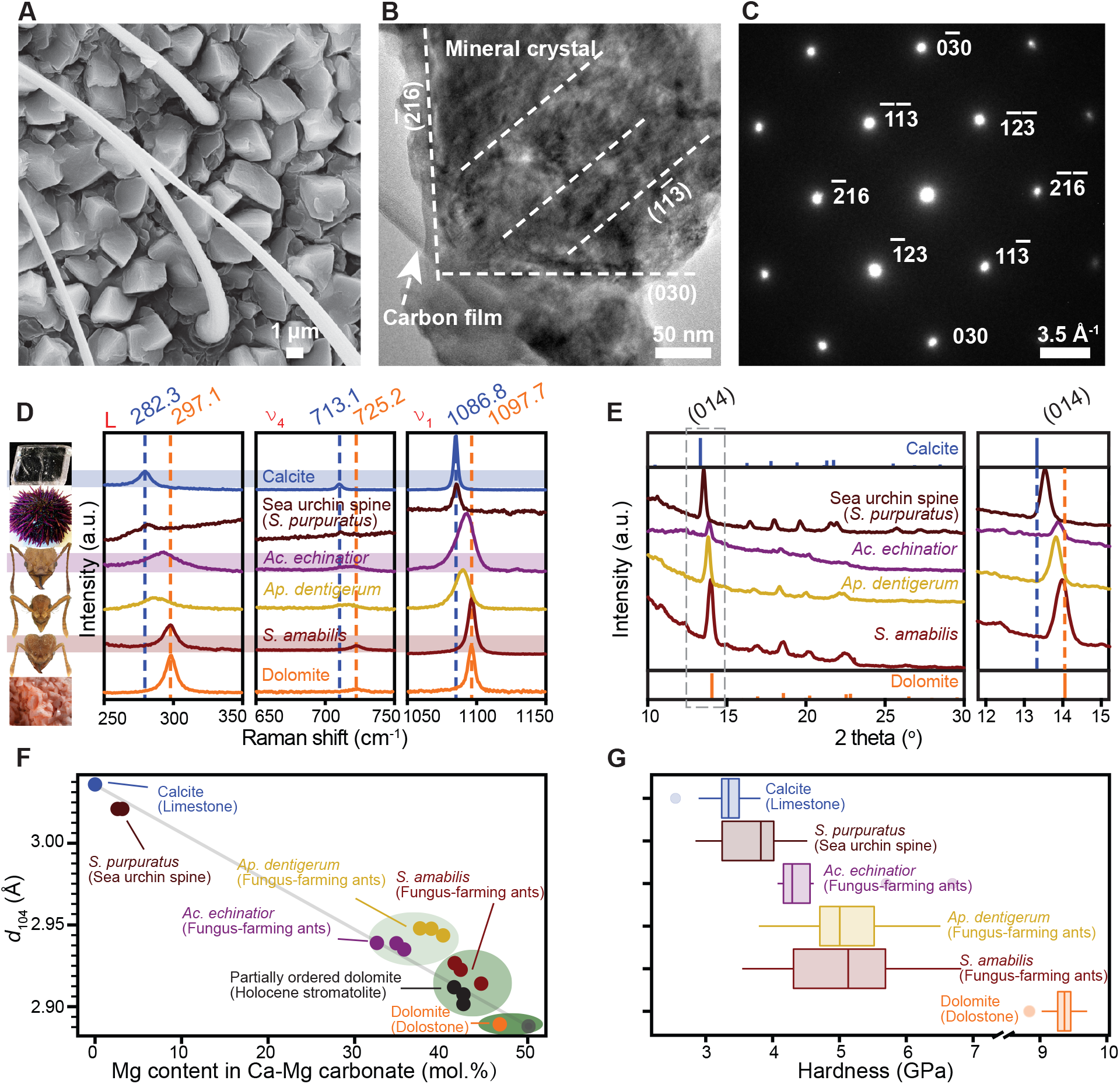
Crystallographic features and mechanical properties of cuticular carbonates in three genera of fungus-growing ants. (**A**) SEM of *S. amabilis* ant cuticle covered in biomineral. **(B)** HRTEM photomicrograph and **(C)** SAED patterns of mineral particle removed from *S. amabilis* ant cuticle. (**D**) Raman spectra and **(E)** powder XRD analysis of various Ca-Mg carbonates from calcite, sea urchin spine, *Ac. echinatior, Ap. dentigerum*, and *S. amabilis* ants. **(F)** MgCO_3_ mol.% versus (104) *d*-spacing value for limestone, sea urchin spine, *Ac. echinatior, Ap. dentigerum, S. amablis*, holocene stromatolite, and dolostone measured by EPMA and XRD. **(G)** Quantitative nano-mechanical properties of various Ca-Mg carbonates from calcite, sea urchin spine, *Ac. echinatior, Ap. dentigerum, S. amabilis* ants, and dolomite, measured by an *in-situ* nanoindenter with a cube-corner probe.

## Discussion

Prior to the discovery of biogenic high-magnesium calcite in the leaf-cutting ant *A. echinatior* (*13*) and to the discoveries reported here in the fungus-farming ants *Sericomyrmex* and *Apterostigma*, high-magnesium calcite had only been documented in a single extant metazoan in the tooth tip of sea urchins (*24*). The incorporation of high concentrations of magnesium into the calcareous biomineral layer in fungus-farming ants correlates with significant increases in armor hardness, presumably providing increased protection. Although magnesium is almost universally present within calcium carbonate, the most widespread biomineral across the animal kingdom, predominantly in the forms of calcite and aragonite, it typically replaces calcium in the crystal lattice only in trace amounts. In aqueous solutions, Mg^2+^ has high affinity with water, forming a strongly bound hydration shell. Carbonates must displace these tightly coordinated water molecules before they can bind Mg^2+^, and this dehydration step creates a kinetic barrier that inhibits the formation of high-magnesium calcites and dolomite (*25*–*27*). The ability of fungus-farming ants to rapidly integrate a substantial amount of magnesium into a calcite lattice is thus remarkable and appears to be unique in terrestrial metazoans.

In the fungus-farming ant *S. amabilis*, we discovered that worker ants possess carbonate biomineral armor with magnesium content reaching as high as 45 mol.%, a level comparable to that found in modern geologic dolomite. Modern geologic dolomite differs from ancient dolomite, formed in the Precambrian and early Paleozoic eras, in the degree of cation ordering. This difference constitutes the ‘dolomite problem’: due to the kinetic barriers imposed by magnesium’s strong hydration shell, modern dolomite forms extremely slowly and laboratory synthesis of dolomite requires high temperatures to overcome these kinetic limitations (*28, 29*). Multiple lines of evidence indicate that the rapidly generated carbonate biomineral in *S. amabilis* ants includes partially ordered dolomite, including: (i) powder XRD patterns indicate a clear smaller (104) *d*-spacing value of 2.923 Å (Fig. 4F) and super-lattice ordering peaks (015), (021), and a slight (101) signal, together comprising the three key cation ordering reflections (Fig. 4E) (*30*); (ii) selected-area electron diffraction (SAED) of individual *S. amabilis* cuticular crystals identified super-lattice reflections (101) and 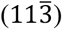 indicating Ca-Mg cation ordering with the alternating layers of calcium and magnesium along the c-axis in the crystal lattice (Fig. 4C) (*21, 22, 31*); and (iii) the Raman spectrum of *S. amabilis* crystals matches that of geologic dolomite rather than of geologic calcite, unlike the crystals in the biomineral layers of the fungus-farming ants *Acromyrmex* and *Apterostigma* (Fig. 4D). The presence of intermediate cation ordering within the crystal lattice in *S. amabilis* is consistent with the theoretical prediction that Ca-Mg ordering starts to occur at the level of 36 mol.% Mg (*22*) and indicates that these ants are able to alter the kinetics of biomineral formation on their cuticles in ways that result not only in increased magnesium integration, but also in the formation of partially ordered dolomite, a biomineral not previously reported from metazoans.

The ability of metazoans to form calcium carbonate biominerals represents a key evolutionary innovation that enabled the diversification of complex body plans and facilitated the colonization of diverse ecological niches. Elucidating the molecular and cellular mechanisms of biomineral formation is essential for understanding evolutionary adaptations, predicting organismal responses to environmental change, and engineering novel biomimetic materials. Calcium carbonate biomineralization in animals is mediated by specialized cells that create controlled microenvironments in which they regulate pH and ion concentrations, including those of calcium and bicarbonate, and in which organic matrix proteins direct crystal nucleation and growth (*32*). Here, we show with ^13^CO_2_-labelling experiments (Fig 3A-3D) that the process of biomineral formation in *S. amabilis* involves the formation of bicarbonate directly on the cuticle via conversion of atmospheric CO_2_. Identifying the proteins involved in the formation of crystals with magnesium content approaching 50% and including some partial ordering could facilitate the design of new biomimetic materials with the favorable properties of increased hardness and stability associated with higher magnesium concentrations.

We have shown that *Sericomyrmex* fungus-farming ants directly convert atmospheric CO_2_ into a carbonate biomineral on the surfaces of their exoskeletons in a process that has parallels with terrestrial geologic weathering (*11, 33*). Geologic carbon mineralization results in the formation of thermodynamically stable, solid carbonate, shaping Earth’s long-term carbon cycle and mediating global climate systems (*34*). In oceans, the abiotic precipitation of stable carbonate minerals involves dissolved inorganic carbon, primarily bicarbonate (HCO_3-_). Additional, biotic carbonate formation is driven by calcareous plankton, which over geological time scales has resulted in the deposition of a CaCO_3_ sedimentary sink that continues to serve as a significant buffer of atmospheric CO_2_ (*35, 36*). In terrestrial environments, carbon mineralization occurs primarily through the weathering of calcium and magnesium silicate minerals, where atmospheric or soil-respired CO_2_ reacts with these minerals to form stable carbonate precipitates in soils and rock formations. Unlike in marine systems, where biogenic calcification is carried out by diverse, widespread, and ecologically dominant organisms, few terrestrial organisms contribute to the formation of carbonates. Terrestrial animals form carbonate structures by combining calcium ions with inorganic carbon dissolved in body fluids, which is then precipitated as calcium carbonate in skeletal elements, shells, or cuticles (*37*–*39*). In contrast, *Sericomyrmex* fungus-farming ants directly convert atmospheric CO_2_ into carbonate biomineral on the surfaces of their exoskeletons.

The benefits of carbonate biosynthesis to fungus-farming ants include the formation of a biomineral body armor capable of resisting infections by microbial pathogens, including entomopathogenic fungi (*13*), and of protecting against attack by other ants, including army ants and specialized agro-predatory ants known to raid their nests to steal their fungus gardens and brood (*40, 41*). Here, we show that the benefits also include the scrubbing of CO_2_ from their subterranean nest cambers. All subterranean-nesting ant colonies face significant challenges with CO_2_ accumulation due to the metabolic activity of workers in confined spaces. Without adequate ventilation, CO_2_ levels can rise to concentrations that would be lethal to most insects. Fungus-farming ants face even greater CO_2_ accumulation challenges due to the combined metabolic activity of dense ant populations and their cultivated fungal gardens (*42, 43*). CO_2_ concentrations can reach levels that reduce the respiration of the symbiotic fungus, potentially compromising colony growth since ant brood feed exclusively on the fungus (*44*). Fungus-farming ants have evolved sophisticated architectural and behavioral solutions to manage nest ventilation. Different nest architectures facilitate varying ventilation strategies: shallow nests with multiple surface connections and thatched nests utilize wind-induced flows and thermal convection, while serial nests with single openings may be constrained by surrounding soil CO_2_ levels (*17*). Colonies of *Sericomyrmex* form serial nests, typically in clay soils that reduce gas exchange (*45*), and as such should experience significant elevated, toxic levels of CO_2_, which they appear to address, at least in part, through carbon sequestration via biomineral formation.

Insights gained from the complex biology of ants have yielded progress towards addressing diverse human challenges, ranging from robotics and network design to sustainability and medicine (*46*–*48*). The highly efficient collective behaviors of ants have inspired algorithms for traffic flow, logistics, and communication networks (*49*–*51*), while research on fungus-farming ant antimicrobial strategies and symbiotic relationships have yielded clues for developing new antibiotics (*52, 53*). *Sericomyrmex* fungus-growing ants engage in biogenic carbon mineralization, directly scrubbing accumulating CO_2_ deep in their subterranean nest chambers and converting it into stable carbonate mineral armor, including dolomite. The challenge presented to the ants by the accumulation of toxic CO_2_ provides a fascinating and instructive natural parallel to human global efforts to mitigate climate change through carbon mineralization, one of the promising strategies being explored for carbon capture and storage, due to the potential for permanently sequestering gigatons of CO_2_ annually through engineered mineral carbonation processes.

## Acknowledgments

We thank J.W. Valley for his valuable review and discussions; Y. Fang for expert help conducting SEM image analysis; U. P. Agarwal and S. A. Ralph for the Raman spectroscopy measurements; J. Morasch for assistance with nanoindentation measurements; J. Sosa-Calvo for ant imaging.

## Author contributions

Conceptualization: H.L., and C.R.C.; Data curation: H.L., Y.F., W.L., J.L., C.-Y.S., X.K., G.B.-M., J.S., X.M., J.-L.H., J.M., L.C., Z.L., T.R.S., R.E.J., and C.R.C.; Funding acquisition: H.L., W.L., G.B.-M., T.R.S., R.E.J., and C.R.C.; Visualization: H.L., Z.L., T.R.S., R.E.J., and C.R.C.; Investigation: H.L., Y.F., W.L., J.L., C.-Y.S., X.K., J.S., X.M., J.-L.H., J.M., L.C., Z.L., T.R.S., R.E.J., and C.R.C.; Writing - original draft: H.L., T.R.S., and C.R.C.; Final version: all authors.

## Data and materials availability

All data is available in the main text or the supplementary materials.

## Supplementary Material

Materials and Methods

Supplementary Text

Figs. S1 to S16

Tables S1 to S5

Movies S1-S2

References (54-74)

